# Increased capillary stalling is associated with endothelial glycocalyx loss in subcortical vascular dementia

**DOI:** 10.1101/2020.04.08.031187

**Authors:** Jin-Hui Yoon, Paul Shin, Jongyoon Joo, Gaon S. Kim, Wang-Yuhl Oh, Yong Jeong

## Abstract

Proper regulation and patency of cerebral microcirculation is crucial for maintaining a healthy brain. Capillary stalling, i.e., the brief interruption of microcirculation mainly by leukocytes, has been observed in several diseases and contributes to disease pathogenesis or progression. However, the underpinning mechanism for leukocyte capillary plugging remains elusive. Therefore, we investigated the mechanism of capillary stalling in mice during the development of subcortical vascular dementia (SVaD), the most common type of vascular dementia characterized by impaired microcirculation and associated pathological features. Longitudinal optical coherence tomography angiography showed increased number of stalled segments as the disease progressed, while two-photon microscopy indicated a less extensive endothelial glycocalyx (EG) in the stalled segments. We also found that increased gliosis and blood-brain barrier leakage were correlated with the increased number of stalled segments. Based on the above, we conclude that EG potentially mediates capillary stalling and can be a therapeutic target of SVaD.

## Introduction

For proper brain functioning, microcirculation is important not only in healthy but also in various pathological conditions^1^. Previous studies on cortical microcirculation have revealed that cerebral capillary flow velocities are heterogeneous, and some portions of capillaries are being frequently stalled^2, 3^. Recent studies have found that capillary stalling occurs naturally and not randomly in multiple capillaries^4, 5^, while another study reported that increased capillary stalling heavily impacts on cerebral blood flow (CBF)^6^. While previous studies have shown that capillary stalling is induced by leukocyte plugging, the location and mechanism of this phenomenon and its consequences still need to be corroborated^6, 7^. The endothelial glycocalyx (EG) is a layer of proteoglycans and glycoproteins at the luminal side of the vessels^8^. This layer has several functions, including the regulation of blood and endothelial cell interactions, such as neutrophil adhesion^9, 10^. Considering its roles and location, the EG is a possible mediator in the mechanism of capillary stalling.

An increased number of stalled segments has been reported in several neurological diseases, such as Alzheimer’s disease and stroke^6, 11^. Here, we investigated capillary stalling in a model of subcortical vascular dementia (SVaD). SVaD is the most frequent type of vascular dementia, impairing microcirculation and appearing as white matter hyperintensity in magnetic resonance imaging^12, 13^. Since the deficits in SVaD develop over time, a longitudinal investigation is required to properly understand the disease. However, no study has examined microcirculation changes, such as alterations in capillary stalling, in SVaD over several months^7, 12^.

In this study, we aimed to elucidate the underpinning mechanism of capillary stalling in SVaD and its correlation with SVaD pathologies. Changes in microcirculation, such as capillary stalling, vessel density, and arterial blood flow, were quantitatively analyzed during SVaD development using longitudinal optical coherence tomography (OCT) angiography (OCTA) and Doppler OCT (D-OCT), which enable label-free visualization of three-dimensional (3D) microvasculature and blood flow^14, 15^. The extent of the EG in the stalled segments was compared with that in the non-stalled segments using two-photon microscopy (TPM). In addition, histopathological analysis was performed to identify the effects of capillary stalling in brain pathology. Our longitudinal imaging study provides integrative understanding on microcirculation changes in normal conditions and in SVaD.

## Results

### Stalled segments are observed in OCT angiograms

Figure 1A shows the experimental timeline of all surgical and imaging procedures. For *in vivo* imaging, a chronic cranial window was implanted on the left somatosensory cortex at 8 weeks of age^16^. Animals were divided into two groups: one used for OCTA and D-OCT imaging (OCTA and D-OCT group: n = 8 for control and n = 9 for SVaD) and the other for OCTA and TPM imaging (OCTA and TPM group, n = 4 for SVaD). In the first group, imaging was performed every week for 9 weeks (W0 ∼ W8) after two weeks of recovery from the window surgery. In the second group, imaging was performed right before the SVaD induction (W0) and 7 weeks after induction (W7). The SVaD was induced at 10 weeks of age using a microcoil at the right common carotid artery (CCA) and an ameroid constrictor at the left CCA (Figure 1B)^17^. After 9 weeks of imaging, the brain was harvested for immunohistochemistry (IHC). A time-averaged OCT angiogram shows the capillary bed with high spatial resolution (Figure 1C). Stalled segments were identified using a binarized capillary map (right panel in Figure 1C) and serial OCT angiograms and are shaded in blue in the depth-projected OCT angiogram shown in Figure 1D. Stalled segments showed a large decrease in the signal at certain time points due to the instant plugging of blood cells (Figure 1D and Supplementary Video 1).

**Figure 1.**
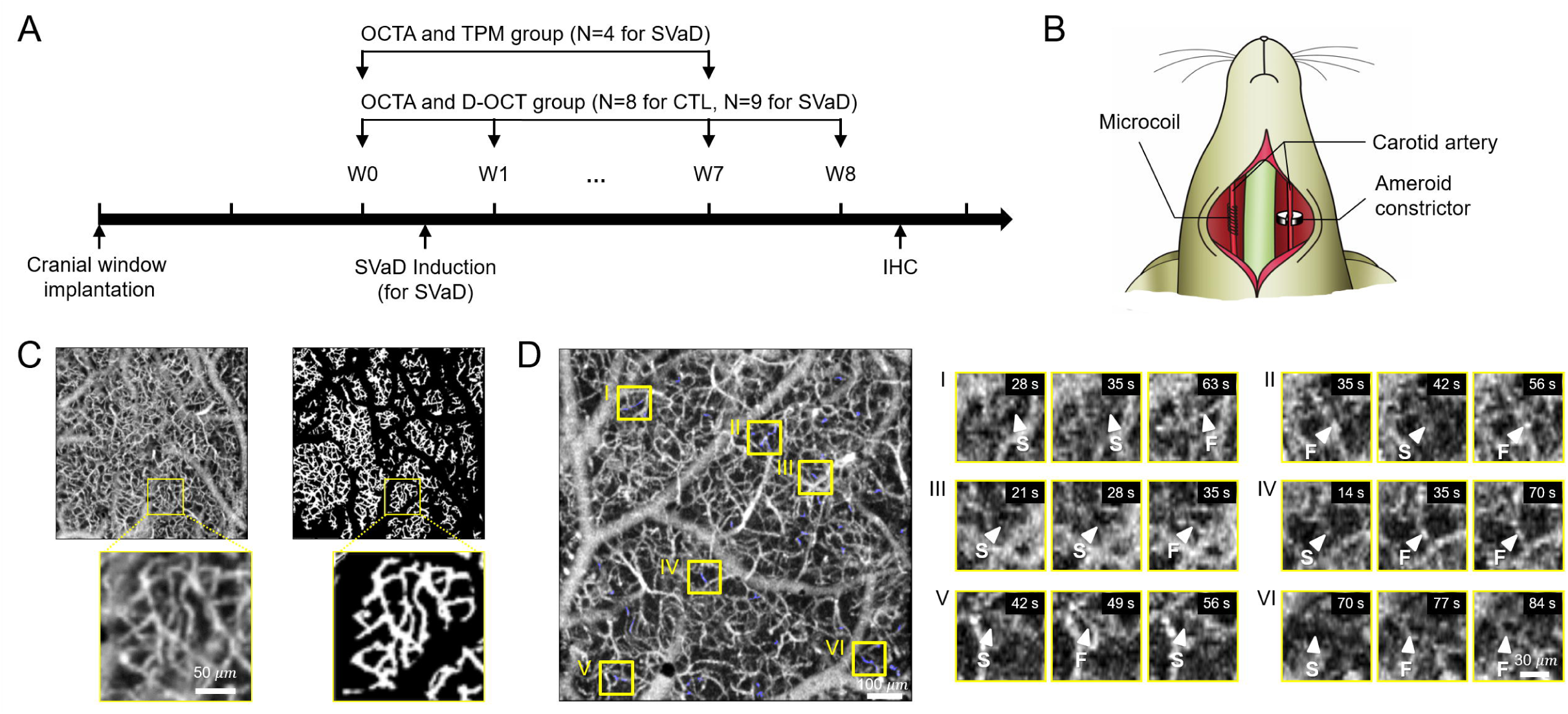
OCTA for the imaging of capillary stalling in SVaD mice. A. Experimental timeline for longitudinal imaging (n = 8 for control and n = 13 for SVaD mice). B. An illustration describing the method for inducing SVaD. An ameroid constrictor was implanted on the left CCA and a microcoil was implanted on the right CCA. C. (left panel) A time-averaged OCT angiogram obtained by averaging all angiograms acquired in 91 seconds. An enlarged image of the region indicated by the yellow box is also shown. (right panel) A binary mask of capillaries obtained using the time-averaged OCT angiogram. The enlarged image of the same region indicated by the yellow box is shown. D. (left panel) A depth-projected *en face* OCT angiogram (150-300 μm from the cortical surface) shows the cerebral microvasculature with high spatial resolution. Positions of stalled segments are shaded in blue. (right panel) Magnified OCT angiograms of the regions indicated by yellow boxes. For each region, capillary stalling appearing at a certain time point is indicated by the letter S, and flowing after or before the stalling is indicated by the letter F. OCTA, optical coherence tomography angiography; OCT, optical coherence tomography; CCA, common carotid artery; SVaD, subcortical vascular dementia.

### Capillary stalling increases with SVaD progression

Alterations in vessel density, defined as the total number of capillary segments in the skeletonized binary vessel mask, as well as density of stalled segments, defined as the ratio of the number of stalled segments to the total number of capillary segments, were measured. Charge-coupled device (CCD) images in Figure 2A show the regions where OCTA was performed in control and SVaD mice. Alterations in microcirculation were hardly detectable in CCD images due to insufficient spatial resolution. Time-averaged OCT angiograms acquired at the same regions during the 9 weeks of imaging showed apparent capillary loss in SVaD mice, while no apparent changes were observed in control mice (Figure 2B). Vessel density (normalized to the baseline value acquired at W0) decreased in SVaD mice (Figure 2C), although not to a significant level (F_(8,104)_ = 1.165, p = 0.3274 for interaction; F_(1,13)_ = 2.058, p = 0.1751 for group factor). Meanwhile, the density and number of stalled segments were significantly higher in SVaD than in control mice (Figure 2D and Supplementary figure 1; F_(8,104)_ = 4.232, p = 0.0002 for interaction; F(1,13) = 7.544, p = 0.0166 for group factor). A sharp increase in the density of stalled segments was observed between one and three weeks after inducing SVaD, possibly associated with the rapid decrease in arterial flow, as depicted in Supplementary Figure 2. At W8, the density of stalled segments in SVaD mice was 2.61 times higher than that in control mice (control: 1.24-fold change, SVaD: 3.24-fold change, p = 0.024). The number of stalled segments at W0 (control: 23.6 ± 2.6, SVaD: 20.9 ± 2.6, p = 0.4939) was much smaller than the total number of capillaries in both control and SVaD mice (control: 4367 ± 379.4, SVaD: 4972 ± 344.5, p = 0.2688). Approximately 40% of the stalled segments identified at W0 were stalled again at least once during the 9 weeks of imaging in both control and SVaD mice (Figure 2E; control: 38.41 ± 4.6, SVaD: 40.77 ± 4.5, p = 0.7298). This result indicates that the stalling did not occur randomly but that certain segments have higher susceptibility to stalling, as mentioned in other studies^4, 5^. It is noteworthy that more than 40% of the re-stalled segments were lost at W8 in SVaD mice, which is twice the rate of capillary segments loss compared to that in control mice (Figure 2F; control: 20.87 ± 3.5, SVaD: 42.89 ± 7.0, p = 0.0042). During the 9 weeks of imaging, there were no differences in physiological signals between control and SvaD mice (Supplementary Figure 3).

**Figure 2.**
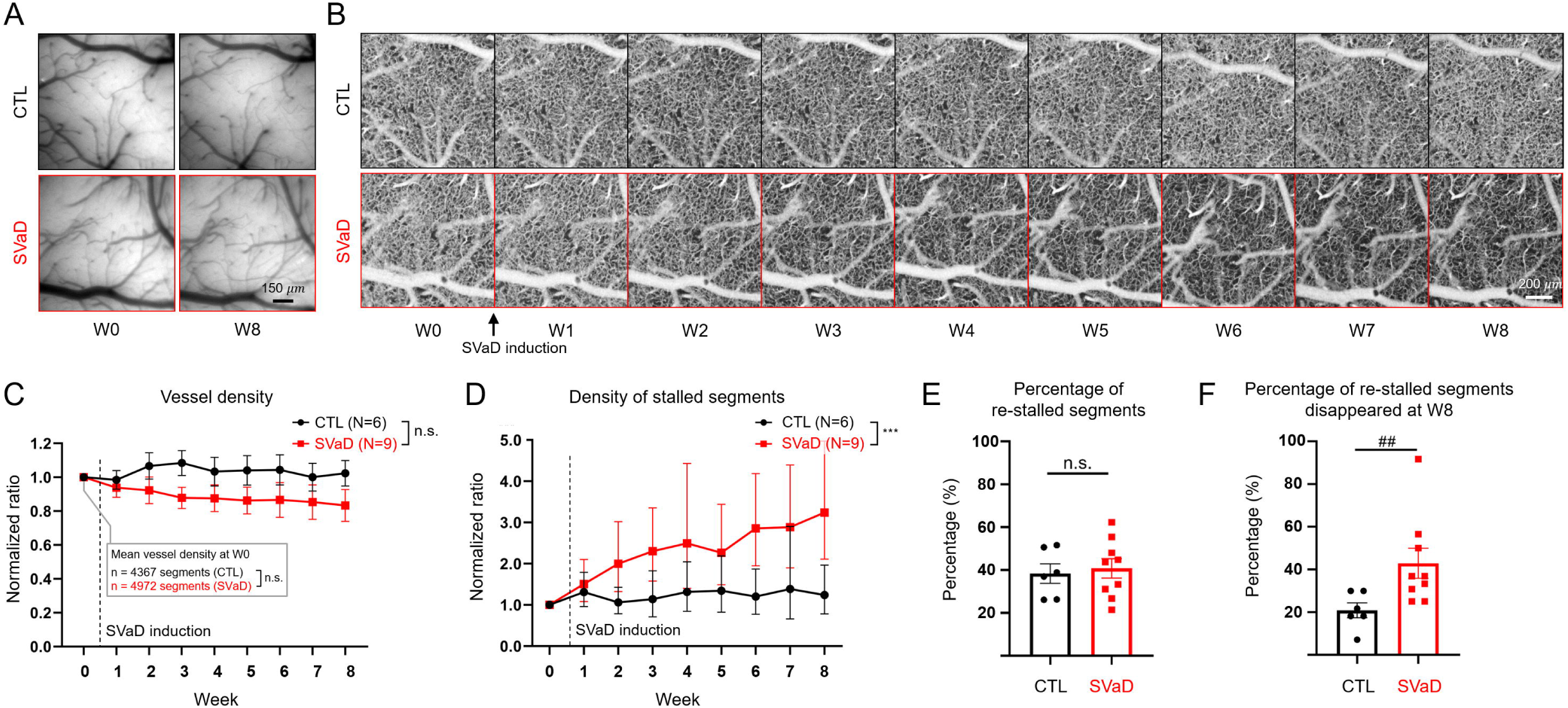
Alterations in microcirculation during disease development. A. CCD images acquired at different time points showing no visible changes in microcirculation both in control and SVaD mice. B. Representative OCT angiograms of control and SVaD mice showing a gradual decrease in vessel density. C. Changes in vessel density in control and SVaD mice. Values are normalized to baseline, and the error bars represent s.e.m. D. Changes in the density of stalled segments in control and SVaD mice. Values are normalized to baseline, and the error bars represent 95% confidence interval of the mean. E. The percentage of re-stalled segments among stalled segments in control and SVaD mice. Data are presented as mean ± s.e.m. F. The percentage of re-stalled segments that disappeared at W8 in control and SVaD mice. Data are presented as mean ± s.e.m. n = 6 for controls and n = 9 for SVaD mice. Two-way repeated measure ANOVA test followed by Bonferroni’s post-hoc test were used (**: p < 0.01, ***: p < 0.001 for the interaction effect of group and time). Log transformation was performed for the density of stalled segments. Two-tailed unpaired student t-test and Mann-Whitney test were used (##: p<0.01 for Mann-Whitney test). CCD, charge-coupled device; OCT, optical coherence tomography; ANOVA, analysis of variance; s.e.m., standard error of the mean; SVaD, subcortical vascular dementia.

### The extent of the endothelial glycocalyx is lower in stalled segments

We investigated the association between capillary stalling and the extent of the EG in SVaD mice. The boundary between endothelial cells and the EG layer was defined as the location where the signal intensity of wheat germ agglutinin lectin (WGA-lectin) staining exceeded the mean plus five times of standard deviation of background noise, while the boundary between the EG layer and blood plasma was defined as the inflection point of the intensity profile of dextran (Figure 3A)^18^. The area under the plot of the WGA-lectin intensity profile between the two boundaries was defined as the extent of the EG. Stalled segments located 150-200 μm from the cortical surface were selected for TPM imaging (shaded in blue in the OCT angiogram in Figure 3B). As shown in a magnified OCT angiogram in Figure 3B, a stalled segments and a nearby non-stalled segments with similar branching order were selected. The area under curve of WGA-lectin signals in the EG layer was generally smaller in the stalled segments than in the non-stalled ones. As depicted in Figure 3C, the extent of the EG was significantly smaller in the stalled segments than in the non-stalled segments both at W0 (non-stalled: n = 17, 42.65 ± 4.957; stalled: n = 10, 22.25 ± 3.029; p = 0.0018, Welch’s t-test) and W7 (non-stalled: n = 17, 32.65 ± 2.921; stalled: n = 11, 15.26 ± 2.298, p = 0.0002, Student t-test).

**Figure 3.**
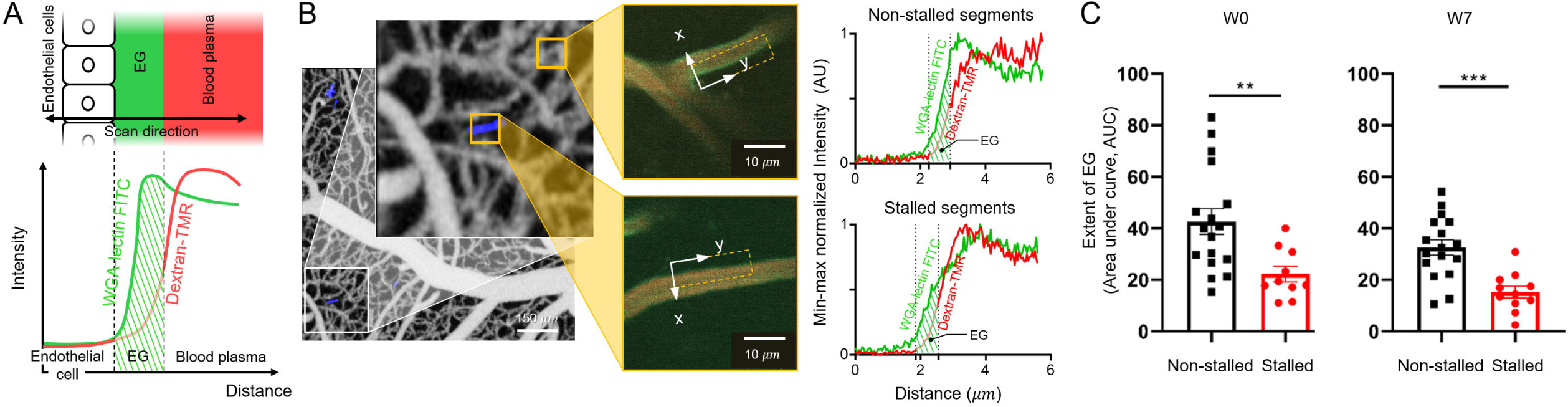
Extent of the EG in stalled segments. A. (top row) An illustration showing the EG layer located at the endothelial surface. (bottom row) The intensity profile of WGA-lectin along the scan direction is shown. The intensity profile of dextran is overlaid. Two vertical dotted lines indicate boundaries among endothelial cells, EG layer, and blood plasma. The region indicated by the diagonal lines is defined as the extent of the EG. B. (left panel) An OCT angiogram with a magnified view of the capillary bed. Stalled segments are shaded in blue. (middle panel) TPM images of selected regions indicated by orange squares in the angiogram showing a non-stalled and a stalled segment, respectively. (right panel) Averaged intensity profiles of WGA-lectin and dextran signals of the capillaries indicated by dotted squares in TPM images, where two vertical dotted lines indicate boundaries between endothelial cells, the EG layer, and blood plasma. Each profile was minimum-maximum normalized for better representation. C. (left panel) The extent of the EG in non-stalled and stalled segments at W0 (n = 17 for non-stalled and n = 10 for stalled segments from four SVaD mice). (right panel) The extent of the EG in non-stalled and stalled segments at W7 (n = 17 for non-stalled and n = 11 for stalled segments from three SVaD mice). Data are presented as mean ± s.e.m. Two tailed unpaired t test was used (**: p < 0.01, ***: p < 0.001, Welch’s t-test and student’s t-test). EG, endothelial glycocalyx; OCT, optical coherence tomography; s.e.m., standard error of the mean; SVaD, subcortical vascular dementia; TPM, two-photon microscopy; WGA-lectin, wheat germ agglutinin lectin.

### Association of reactive astrocytes and blood-brain barrier (BBB) leakage with capillary stalling in SVaD

We observed reactive astrocytes stained with glial fibrillary acidic protein (GFAP) (left column) and found they were increased in SVaD mice (Figure 4A, magenta color). In addition, we observed increased BBB leakage in SVaD mice, represented by endogenous immunoglobulin G (IgG) leakage (right column) (orange color). In SVaD mice, penetrating arterioles and capillaries were ensheathed by reactive astrocyte endfeet (Figure 4B, left and middle panels). In control mice, reactive astrocytes and vessels with reactive astrocyte endfeet were rarely observed (Supplementary Figure 4). As shown in Figure 4B (right panel), endogenous IgG signal was highly accumulated around capillaries, as pointed by white arrowheads. IHC analysis was performed using the 5 control and 9 SVaD mice that had been used for the OCTA analysis. Figure 4C shows the significantly higher GFAP expression (control: 2.505 ± 0.749, SVaD: 8.018 ± 1.061, student’s t test p = 0.0039) and IgG deposition (control: 326.2 ± 48.99, SVaD: 523.8 ± 29.51, Mann-Whitney test p = 0.007) in SVaD than in control mice, in agreement with previous observations in other SVaD models^12, 17^. To examine the possible correlations between capillary stalling and the histopathological features of SVaD, we measured the correlation between the severity of capillary stalling and GFAP (or IgG) expression. Both gliosis and BBB leakage showed positive correlations with the severity of capillary stalling (gliosis: r = 0.5367, p = 0.0479; BBB leakage: r = 0.5557, p = 0.0391). No significant correlation was observed within each separate group.

**Figure 4.**
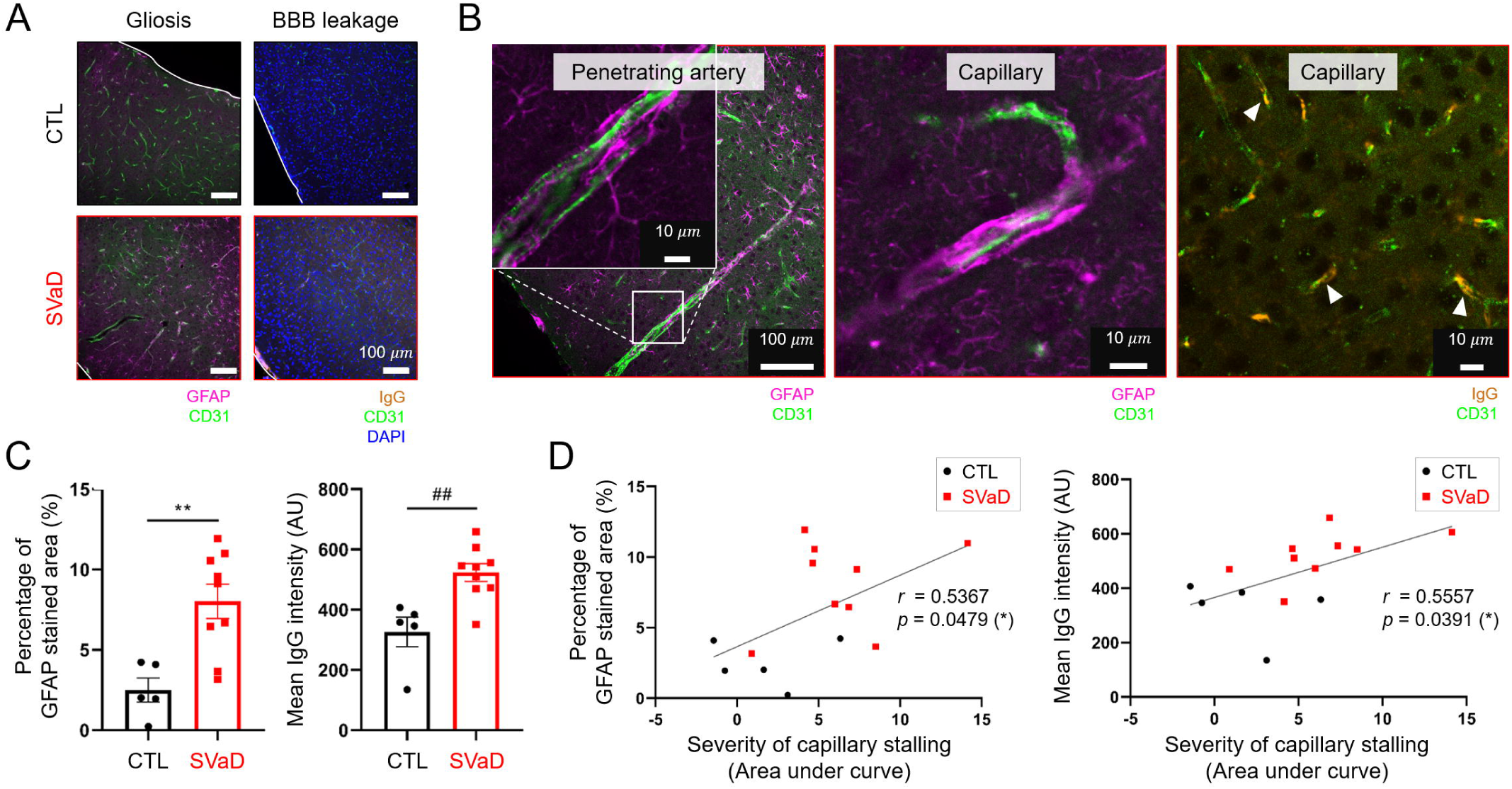
Correlation between capillary stalling and histopathological features of SVaD. A. (left column) IHC images showing the expression of GFAP due to gliosis (indicated in magenta color). (right column) IHC images showing endogenous IgG leakage due to the BBB disruption (indicated in orange). The white solid line in each IHC image indicates the cortical surface. B. IHC images of an SVaD mouse. (left panel) A penetrating artery ensheathed by reactive astrocyte endfeet. An enlarged image of the corresponding artery is shown. (middle panel) A capillary ensheathed by reactive astrocyte endfeet. (right panel) The accumulation of leaked endogenous IgG around capillaries, pointed by white arrowheads. C. The degrees of GFAP and IgG expression in control (n = 5) and SVaD (n = 9) mice. D. (left panel) A scatter-plot between the severity of capillary stalling and GFAP expression. (right panel) A scatter-plot between the severity of capillary stalling and endogenous IgG expression. Control mice (n = 5) are indicated by black square dots and SVaD mice (n = 9) are indicated by red dots. Data are presented as mean ± s.e.m. Two tailed unpaired t test and Mann-Whitney test were used (**: p < 0.01, ##: p < 0.01 for student t-test and Mann-Whitney test, respectively). Pearson coefficient was calculated (*: p < 0.05). IHC, immunohistochemistry; BBB, blood-brain barrier; GFAP, glial fibrillary acidic protein; s.e.m., standard error of the mean; SVaD, subcortical vascular dementia.

## Discussion

We investigated the longitudinal alterations in microcirculation in a SVaD model using OCT. Twice or more as many stalled capillary segments were observed in SVaD than in control mice, while more than 40% of the re-stalled segments disappeared at the final week of imaging in the former, which was twice as many segments disappearing in control mice. The extent of the EG in the stalled segments was significantly smaller than that in the non-stalled segments. Further, capillary stalling correlated with gliosis and BBB leakage.

SVaD show various pathological features of microcirculation impairments. The gradual decrease in vessel density is one of the typical features^19, 20^. By utilizing OCTA, we demonstrated that the vessel density decreased noticeably more in some of the SVaD than in control mice during the 9 weeks of imaging, although the overall difference was not statistically significant. Notably, the density of stalled segments began to increase with the induction of SVaD and became 2.61 times higher than that in control mice after 8 weeks. Capillary stalling has been identified not only during disease progression but also before induction of the disease^4, 5^. To examine whether stalling occurred randomly or preferentially to certain segments, we repeatedly imaged the stalled segments for 9 weeks. Both in control and SVaD mice, about 40% of the stalled segments at baseline were stalled again at least once during that period, implying that certain capillary segments are more susceptible to stalling than others, although we found no difference between the two groups. This finding indicates that capillary stalling is part of a physiologic function, although the exact nature of which remains elusive. However, among the re-stalled segments, the proportion of segments that disappeared at the final week of the imaging (W8) was approximately twice as large in SVaD (40%) than in control (20%) mice, suggesting that the induction of SVaD promotes a more rapid loss of the cerebral microvasculature.

Considering the limited acquisition time, there must be missed portion of stalled segments that was not detected. Furthermore, capillary segments that stalled for a shorter time than the temporal resolution limit of the imaging modality (7 seconds) would be also missed. Despite the likelihood of underestimating the number of stalled segments, our results clearly show the rapid progression of capillary stalling in SVaD.

Recent studies have revealed the significance of capillary stalling in aging, stroke, Alzheimer’s disease, polycythemia and thrombocythemia^3, 4, 5, 6, 11, 21, 22, 23^. In fact, capillary stalling largely impacts on CBF, and cognitive functions can be improved by suppressing capillary stalling in Alzheimer’s disease^6, 24^. The underlying mechanism of capillary stalling involves various intravascular components, such as blood clots, red blood cells, and leukocytes. Using a cerebral hypoperfusion model, a previous study demonstrated that capillary stalling is generated by the plugging of leukocytes and not by platelet activation^7^. In another study, leukocyte plugging in stalled capillaries was also identified in an Alzheimer’s disease model, demonstrating that it is a plausible mechanism underlying capillary stalling^6, 7^. According to our results, a large increase in the number of stalled segments was observed during the first 4 weeks after SVaD induction, when the arterial flow speed rapidly decreased. This result supports previous observations regarding the close relationship of the incidence of capillary stalling to the overall flow supplied to the vessels^7^. However, the reasons for which the plugging of blood cells occurs still remain unclear. Structural characteristics of the vessels such as vessel length and tortuosity could also be associated with capillary stalling, although a previous study showed no such correlation^4^.

In this study, we considered the EG as a possible candidate involved in the plugging of blood cells, considering it is the interface structure between endothelial cells and the blood stream. Utilizing TPM, we measured the extent of the EG in the stalled segments and compared it to that in the non-stalled segments. Our results showed that the EG extent was significantly smaller in the stalled than in the non-stalled segments. This was observed not only in the SVaD but also in normal mice. The EG has several functions, including the regulation of immune cell interactions with the endothelium, transmission of fluid shear stress, and control of substance transport^8, 25, 26^. The EG layer of brain capillaries has a thickness of typically around 400 nm^18^. Several hypotheses have been suggested to explain the interaction between the EG and neutrophils^26^. One of the convincing hypotheses proposes that the neutrophil-endothelial adhesion may be prevented by the charged repulsion between the negatively-charged leukocyte surface layer and the negatively-charged EG layer, which covers adhesion molecules such as intercellular adhesion molecule 1^8^. Shedding of the EG reduces charge repulsion and makes the EG layer easy to be compressed by leukocytes, thereby facilitating leukocyte-endothelial adhesion^26^. Another previous study showed enhanced leukocyte adhesion in the retina of knockout mice for syndecan 1, a component of the EG^27^. In addition, several studies have shown degradation of the EG in diseases characterized by decreased CBF, such as malaria infection, multiple sclerosis, and stroke^28, 29, 30^. Based on these previous studies and our results, we suggest that the loss of the EG in capillaries, caused by the pathological progression of SVaD, may induce greater leukocyte adhesion to the vascular endothelium and, thereby, promote capillary stalling. In the SVaD model used in this study, the gradual decrease in arterial flow speed and pulsatility index, which is influenced by artery inflow and vessel resistance, was followed by impaired microcirculation (Supplementary Figure 2)^31, 32^. Since the production and maintenance of the EG closely correlates with fluid shear stress, it is plausible that the decreased capillary flow speed may be a primary factor affecting the EG loss^25, 33, 34^. Furthermore, this correlation could explain why, even in the normal brain, the EG of some capillaries appeared to have a smaller extent. This can be attributed to slower red blood cell and plasma velocity in the stalled segments^4^. Thus, it is clear that the EG is deeply involved in capillary stalling both under normal and pathologic conditions.

Further, we examined the consequence of increased capillary stalling. We considered that alterations in the microcirculation, such as due to stalling, would reduce the efficiency of oxygen delivery to tissues that are presumably associated with the SVaD pathogenesis^21, 35^. We thus investigated the relationship of stalling with SVaD pathological findings, such as gliosis and BBB integrity impairment. We observed significantly more reactive astrocytes and a higher degree of BBB leakage in SVaD mice, in agreement with the findings of previous studies^12, 17, 36^. Local hypoxia by stalling may contribute to BBB leakage or increased gliosis^37, 38^. Oppositely, both gliosis and BBB leakage could affect the proper regulation of capillary flow during SVaD development, since astrocytes are mainly involved in the regulation of microcirculation^39, 40, 41, 42^, and BBB dysfunction can cause immune cells to enter the brain^43^. Regardless, we suggest that capillary stalling is associated with the presence of reactive astrocytes and BBB leakage. Taken together, our result suggest that under decreased arterial blood flow in SVaD, reduced shear stress induces loss of the EG in capillaries that may cause increased capillary stalling by leukocyte plugging. In turn, this causes gliosis and BBB disruption, which may lead to a vicious cycle leading to additional capillary stalling.

The results of the study should be viewed in light of some limitations. In this study, OCT and TPM imaging were only performed in the cortex of mice due to the relatively shallow imaging depth of these modalities. Notably, the effect of isoflurane should also be considered, as it may induce capillary stalling, as previously reported^4, 44^. Nevertheless, our results clearly show the significant alterations in the microcirculation of SVaD mice compared to control.

In conclusion, the EG plays a critical role in the mechanism of capillary stalling and associated pathologies in SVaD. We expect that this study will lead to a deeper understanding of the alterations in microcirculation and therapeutic targets in SVaD.

## Material and Methods

### Animal preparation and imaging procedures

All animal preparations and imaging procedures were approved by the Institutional Animal Care and Use Committee (IACUC) of Korea Advanced Institute of Science and Technology (KA2018-04) and were performed in accordance with ARRIVE (Animal Research: Reporting in vivo Experiments) guidelines (https://www.nc3rs.org.uk/arrive-guidelines). Male C57BL/6J mice were used for the study. To implant a chronic cranial window, mice were anesthetized with mixture of Zoletil (30 mg/kg) and Xylazine (10mg/kg) via intraperitoneal injection and a cover glass and head-plate were implanted on the left somatosensory cortex at 8 weeks of age as previously described^16^. To induce SVaD, mice were anesthetized with mixture of Zoletil (30 mg/kg) and Xylazine (10 mg/kg) via intraperitoneal injection and placed on a heat blanket (55-7020, Harvard Apparatus) with temperature feedback via rectal probe to maintain body temperature at 37 °C. A spiral microcoil with an internal diameter of 0.18 mm (Sawane Spring Co.) was placed at the right CCA, and an ameroid constrictor with an inner diameter of 0.5 mm (Research Instrument SW) was placed at the left CCA, as described previously ^17^. Sham surgery was performed for control mice. The brain was collected for IHC after the end of imaging. Throughout imaging, animals were anesthetized with isoflurane (5% for induction, and 1.5∼1.7% for maintenance with a gas mixture of 40% oxygen and 60% air), and physiological signals, such as SO_2_, heart rate, and body temperature were monitored (Physio-Suite, Kent Scientific).

### OCTA and D-OCT imaging

A swept-source OCT system with a center wavelength of 1.3 μm and an A-scan rate of 240 kHz was used, as previously described ^45^. OCTA was performed with a field of view of 1 mm × 1mm. In 3D OCT angiograms, maximum projection was performed in a depth direction over two depth intervals, 0-150 μm and 150-300 μm from the cortical surface. Since most large vessels are located at the surface, and capillaries are embedded deeper in the tissue, the focus of the OCT beam was positioned approximately at 220 μm from the surface. Between the two *en face* OCT angiograms, the one with the deeper depth interval was used for analysis. Serial OCT angiograms were continuously acquired every 7 seconds for 91 seconds to observe temporal changes in microcirculation. Each angiogram consisted of 1000 A-scans (in the x-direction) and 200 B-scans (in the y-direction). The axial and lateral resolution of the system was 10 μm. In OCTA, the inter B-scan time interval is a critical factor that determines the slowest detectable flow ^45, 46^. In this study, the inter B-scan time interval was set to 4.2 milliseconds, which is sufficient to increase the signals of most capillaries to the saturation limit. After OCTA, D-OCT imaging was performed to measure the arterial flow. A single pial arteriole directly feeding the region where OCTA was performed was selected. The vessel was repeatedly scanned across its width along a 250-μm line. The corresponding D-OCT imaging provides a quantitative measurement of the two-dimensional flow profile of the pial artery^47^. Imaging was performed for 30 seconds at a temporal resolution of 0.06 seconds. The number of A-scans per each cross-sectional D-OCT image was 5000.

### OCTA and TPM imaging

A temporal series of OCT angiograms of the mouse cortex was obtained before TPM imaging, as described above. Stalled segments located 150-200 μm from the cortical surface were identified using 3D OCT angiograms and used for EG imaging. Capillaries close to the surface were selected to identify them easily later for TPM imaging. Each animal was moved to the TPM device (LSM 510, Zeiss) after the OCTA. A femtosecond laser with a center wavelength of 800 nm (Chameleon Ultra™, Coherent Inc.) and a Plan-Apochromat 20x/1.0 DIC VIS-IR water immersion objective lens were used for imaging. WGA-lectin conjugated with fluorescein isothiocynate (WGA-lectin FITC) was dissolved in saline (1mg/mL) and injected intravenously via the tail vein 45 minutes before imaging at a dose of 6.25mg/kg, to label the EG, as described previously^18^. Dextran (2 MDa; 0.25% (w/v) in saline) conjugated with tetramethylrhodamine (Dextran-TMR; Thermo Fisher) was also intravenously injected just before the imaging at a volume of 100 μL to label blood plasma. Stalled segments found in OCT angiograms were two-dimensionally scanned one by one by TPM. The field of view was 60 μm^2^, with pixel resolution of 1024 × 1024. The pixel dwell time was 0.64 μs, and each line was generated by averaging 8 lines. The TPM was performed within 1 hour to avoid signal reduction.

### Immunohistochemistry (IHC)

Animals were deeply anesthetized by intraperitoneal injection of ketamine (100 mg/kg) and xylazine (10 mg/kg), and phosphate-buffered saline (PBS) was perfused transcardially. After PBS perfusion, brains were fixed with transcardial perfusion of 4% paraformaldehyde. For immunofluorescence staining, brains were submerged in 4% paraformaldehyde 12 h and transferred to 30 % (w/v) sucrose solution for cryoprotection. Samples were embedded in optimal cutting temperature compound and sliced coronally at a 30-μm thickness. All coronal slices were obtained at the region where OCT imaging was performed. Brain slices were blocked in 5 % normal donkey serum dissolved in PBS and 0.01 % Triton X-100 (blocking solution) after three times of PBS washing. The primary antibodies anti-PECAM-1 (1:1000; MAB1398Z, EMD Millipore) and anti-GFAP (1:1000; ab4674, Abcam) were used to tag endothelial cells and reactive astrocytes, respectively. Secondary antibodies with conjugated fluorophores, anti-Armenian hamster antibody with Alexa Fluor 488 (1:1000; 127-545-160, Jackson ImmunoResearch) and anti-chicken antibody with Alexa Fluor 647 (1:1000; ab150175, Abcam), were used for confocal imaging. To detect BBB leakage, endogenous IgG staining was performed using anti-mouse IgG with Alexa Fluor 594 (1:1000; ab150116, Abcam) without a primary antibody. Primary and secondary antibodies were dissolved in blocking solution. Brain slices were incubated in primary antibody solution at 4 °C overnight, followed by secondary antibody solution incubation at 4 °C overnight. Brain slices were mounted using Vectashield mounting medium (H-1200, Vector Laboratories). Confocal microscopy (Nikon C2, Nikon) was utilized for imaging all brain slices, with a Plan Apo 20x/0.75 objective lens and a field of view of 635 μm x 635 μm. Imaging was performed to cover the region where OCT angiograms were obtained, i.e., the region located 0-300 μm away from the cortical surface.

### Data analysis

Changes in capillary stalling and vessel density were measured using *en face* OCT angiograms. A time-averaged OCT angiogram was generated by repeatedly averaging all angiograms acquired within 91 seconds. The overall contrast of the averaged OCT angiogram was adjusted by contrast limited histogram equalization. After equalization, vessels were segmented using the Weka trainable segmentation plugin in Fiji software ^48, 49^. Shadows from large pial vessels were removed from the image. Following the above procedures, a binary mask of the capillary network was obtained, which was then skeletonized. Vessel density was defined as the total number of skeleton segments. Capillary stalling was evaluated using a temporal series of OCT angiograms. Each angiogram was multiplied by the binary mask to clearly identify the vessels. Capillaries that showed a sudden drop in the signal at least once during the 91 seconds of acquisition were defined as stalled segments and were identified automatically using MATLAB (Mathworks), and double checked by manual inspection. The density of stalled segments was defined as the number of stalled segments divided by vessel density to correct for possibly high stalled segments number due to high vessel density. A time course of changes in vessel density, and the density of stalled segments during 9 weeks of imaging were obtained for each animal. Each time course was normalized to the baseline course (W0), and the area under the curve of the time course of the density of stalled segments was defined as the severity of capillary stalling. Some portions of stalled segments at baseline (W0) were stalled again during 9 weeks of imaging; these capillaries were defined as re-stalled segments. Alterations in vessel density and the density of stalled segments were investigated using 6 control and 9 SVaD mice. Two of 8 control mice were excluded from analysis due to the regrowth of the thick dura mater in the imaging region, which affected the quality of the OCT angiograms.

Arterial hemodynamic parameters, such as vessel diameter, flow speed, and pulsatility were measured using D-OCT imaging. Procedures for measuring the diameter and flow speed were described in our previous work ^45^. Pulsatility was calculated using Gosling’s pulsatility index (PI) equation ^50^. Among the 9 SVaD mice, one was excluded from analysis, as the flow of the pial artery decreased below the minimum detectable limit of D-OCT measurement before W8. In addition, two SVaD mice were excluded from the pulsatility analysis because their heat beat frequencies were too high compared to the frame rate of the D-OCT imaging for more than 4 imaging sessions.

TPM images were analyzed using Fiji software. The intensity profiles of WGA-lectin FITC and dextran-TMR in the direction perpendicular to the vessel wall were measured, as previously described ^18^. The boundary between the EG layer and blood plasma was defined as the inflection point of the minimum-maximum normalized intensity profile of dextran-TMR, while the boundary between endothelial cells and EG the layer was defined as the location where the intensity of WGA-lectin exceeded the mean plus five times the standard deviation of background noise. The area under curve of the intensity profile of WGA-lectin FITC between the boundaries was defined as the extent of EG. TPM image analysis was performed using 4 SVaD mice. At W8, three out of 4 SVaD mice were used for imaging, as one mouse died before W8.

For IHC data analysis, all confocal images were analyzed using Fiji software. To evaluate gliosis, the percentage of GFAP-positive pixels among all pixels within the region of interest was measured. BBB disruption was evaluated as the mean fluorescence intensity of the anti-IgG antibody signal in the image. IHC analysis was performed using the 5 control and 9 SVaD mice that were used for the OCTA analysis. One of the 6 control mice was excluded from the IHC analysis due to severe brain damage that occurred during brain harvesting.

### Statistical analysis

Statistical analysis was performed using GraphPad Prism 8 (GraphPad Software) and SPSS 25 (IBM). All data are presented as mean ± standard error of the mean (s.e.m.) or mean ± 95% confidence interval. Data were tested for normality using Kolmogorov-Smirnov and Shapiro-Wilk tests. Differences between groups were tested using parametric or non-parametric statistics depending on the distribution of data. For comparing two groups, student’s t-test, Welch’s t-test or Mann-Whitney test were used. All longitudinal data from OCT imaging were tested using two-way repeated measure analysis of variance, followed by Bonferroni’s correction. A log-transformation (natural logarithm) was performed for the density of stalled segments, number of stalled segments, and arterial flow speed data to satisfy normality. For examining the correlation between parameters, Pearson correlation coefficient was calculated. Details regarding the number of samples, p values, and other notes for each group are included in the Figures or figure legends.

## Acknowledgments

This research was supported by the Brain Research Program through the National Research Foundation of Korea (NRF), funded by the Ministry of Science and ICT (2016M3C7A1913844) (W.-Y. O. and Y. J.).

## Data availability

The datasets generated during and/or analyzed during the current study are available from the corresponding author on reasonable request.

## References

1. Y. Itoh, N. Suzuki, Control of brain capillary blood flow. J. Cereb. Blood. Flow. Metab. 32, 1167–1176 (2012).

2. D. Kleinfeld, P.P. Mitra, F. Helmchen, W. Denk, Fluctuations and stimulus-induced changes in blood flow observed in individual capillaries in layers 2 through 4 of rat neocortex. Proc. Natl. Acad. Sci. U. S. A. 95, 15741–15746 (1998).

3. A. Villringer, A. Them, U. Lindauer, K. Einhaupl, U. Dirnagl, Capillary perfusion of the rat brain cortex. An in vivo confocal microscopy study. Circ. Res. 75, 55–62 (1994).

4. S.E. Erdener, et al., Spatio-temporal dynamics of cerebral capillary segments with stalling red blood cells. J. Cereb. Blood. Flow. Metab. 39, 886–900 (2019).

5. P. Reeson, K. Choi, C.E. Brown, VEGF signaling regulates the fate of obstructed capillaries in mouse cortex. Elife 7, e33670 (2018).

6. J.C. Cruz Hernandez, et al., Neutrophil adhesion in brain capillaries reduces cortical blood flow and impairs memory function in Alzheimer’s disease mouse models. Nat. Neurosci. 22, 413–420 (2019).

7. K. Yata, et al., In vivo imaging of the mouse neurovascular unit under chronic cerebral hypoperfusion. Stroke. 45, 3698–3703 (2014).

8. S. Reitsma, D.W. Slaaf, H. Vink, M.A. van Zandvoort, M.G. oude Egbrink, The endothelial glycocalyx: composition, functions, and visualization. Pflugers. Arch. 454, 345–359 (2007).

9. S. Massena, et al., A chemotactic gradient sequestered on endothelial heparan sulfate induces directional intraluminal crawling of neutrophils. Blood. 116, 1924–1931 (2010).

10. E.P. Schmidt, et al., The pulmonary endothelial glycocalyx regulates neutrophil adhesion and lung injury during experimental sepsis. Nat. Med. 18, 1217–1223 (2012).

11. S.E. Erdener, et al., Dyanmic capillary stalls in reperfused ischemic penumbra contribute to injury: A hyperacute role for neutrophils in persistent traffic jams. J. Cereb. Blood. Flow. Metab., 271678X20914179 (2020)

12. E.-S. Lee, J.-H. Yoon, J. Choi, F.R. Andika, T. Lee, Y. Jeong, A mouse model of subcortical vascular dementia reflecting degeneration of cerebral white matter and microcirculation. J. Cereb. Blood. Flow. Metab. 39, 44–57 (2019).

13. J.H. Lee, et al., Identification of pure subcortical vascular dementia using 11C-Pittsburgh compound B. Neurology. 77, 18–25 (2011).

14. D. Huang, et al., Optical coherence tomography. Science. 254, 1178–1181 (1991).

15. H. Radhakrishnan, V.J. Srinivasan, Compartment-resolved imaging of cortical functional hyperemia with OCT angiography. Biomed. Opt. Express. 4, 1255–1268 (2013).

16. G.J. Goldey, et al., Removable cranial windows for long-term imaging in awake mice. Nat. Protoc. 9, 2515–2538 (2014).

17. Y. Hattori, et al., A novel mouse model of subcortical infarcts with dementia. J. Neurosci. 35, 3915–3928 (2015).

18. J.-H. Yoon, E.-S. Lee, Y. Jeong, In vivo Imaging of the Cerebral Endothelial Glycocalyx in Mice. J. Vasc. Res. 54, 59–67 (2017).

19. C. Iadecola, The pathobiology of vascular dementia. Neuron. 80, 844–866 (2013).

20. J. Alber, et al., White matter hyperintensities in vascular contributions to cognitive impairment and dementia (VCID): Knowledge gaps and opportunities. Alzheimers. Dement. (N Y). 5, 107–117 (2019).

21. S.E. Erdener, T. Dalkara, Small vessels are a big problem in neurodegeneration and neuroprotection. Front. Neurol. 10, 889 (2019).

22. T.P. Santisakultarm, et al. Stalled cerebral capillary blood flow in mouse models of essential thrombocythemia and polycythemia vera revealed by in vivo two-photon imaging. J. Thromb. Haemost. 12, 2120–2130 (2014).

23. B. Schager, C.E. Brown, Susceptibility to capillary plugging can predict brain region specific vessel loss with aging. J. Cereb. Blood. Flow. Metab. 271678X19895245 (2020).

24. O. Bracko, B.N. Njiru, M. Swallow, M. Ali, M. Haft-Javaherian, C.B. Schaffer, Increasing cerebral blood flow improves cognition into late stages in Alzheimer’s disease mice. J. Cereb. Blood. Flow. Metab. 271678X19873658 (2019).

25. J.M. Tarbell, M.Y. Pahakis, Mechanotransduction and the glycocalyx. J. Intern. Med. 259, 339–350 (2006).

26. A. Marki, J.D. Esko, A.R. Pries, K. Ley, Role of the endothelial surface layer in neutrophil recruitment. J. Leukoc. Biol. 98, 503–515 (2015).

27. M. Gotte, et al., Role of syndecan-1 in leukocyte-endothelial interactions in the ocular vasculature. Invest. Ophthalmol. Vis. Sci. 43, 1135–1141 (2002).

28. B. DellaValle, H. Hasseldam, F.F. Johansen, H.K. Iversen, J. Rungby, C. Hempel, Multiple soluble components of the glycocalyx are increased in patient plasma after ischemic stroke. Stroke. 50, 2948–2951 (2019).

29. B. DellaValle, et al., Detection of glycan shedding in the blood: new class of multiple sclerosis biomarkers? Front. Immunol. 9, 1254 (2018).

30. C. Hempel, J. Sporring, J.A.L. Kurtzhals, Experimental cerebral malaria is associated with profound loss of both glycan and protein components of the endothelial glycocalyx. FASEB J. 33, 2058–2071 (2019).

31. D.H. Evans, W.W. Barrie, M.J. Asher, S. Bentley, P.R. Bell, The relationship between ultrasonic pulsatility index and proximal arterial stenosis in a canine model. Circ. Res. 46, 470–475 (1980).

32. K. Washida, Y. Hattori, M. Ihara, Animal models of chronic cerebral hypoperfusion: from mouse to primate. Int. J. Mol. Sci. 20, 6176 (2019).

33. Y. Zeng, J.M. Tarbell, The adaptive remodeling of endothelial glycocalyx in response to fluid shear stress. PLoS ONE. 9, e86249 (2014).

34. G. Wang, et al., Shear stress regulation of endothelial glycocalyx structure is determined by glucobiosynthesis. Arterioscler. Thromb. Vasc. Biol. 40, 350–364 (2020).

35. L. Ostergaard, et al., Cerebral small vessel disease: Capillary pathways to stroke and cognitive decline. J. Cereb. Blood. Flow. Metab. 36, 302–325 (2016).

36. A.H. Hainsworth, H.S. Markus, Do in vivo experimental models reflect human cerebral small vessel disease? A systematic review. J. Cereb. Blood. Flow. Metab. 28, 1877–1891 (2008).

37. J.I. Na, et al., The HIF-1 inhibitor YC-1 decreases reactive astrocyte formation in a rodent ischemia model. Am. J. Transl. Res. 7, 751–760 (2015).

38. P. Venkat, M. Chopp, J. Chen, Blood-brain barrier disruption, vascular Impairment, and ischemia/reperfusion damage in diabetic stroke. J. Am. Heart. Assoc. 6, e005819 (2017).

39. H. Girouard, C. Iadecola, Neurovascular coupling in the normal brain and in hypertension, stroke, and Alzheimer disease. J. Appl. Physiol. (1985). 100, 328–335 (2006).

40. C.N. Hall, et al., Capillary pericytes regulate cerebral blood flow in health and disease. Nature. 508, 55–60 (2014).

41. A. Mishra, Binaural blood flow control by astrocytes: listening to synapses and the vasculature. J. Physiol. 595, 1885–1902 (2017).

42. A. Mishra, J.P. Reynolds, Y. Chen, A.V. Gourine, D.A. Rusakov, D. Attwell, Astrocytes mediate neurovascular signaling to capillary pericytes but not to arterioles. Nat. Neurosci. 19, 1619–1627 (2016).

43. R. Daneman, A. Prat, The blood-brain barrier. Cold. Spring. Harb. Perspect. Biol. 7, a020412 (2015).

44. A.M. Slupe, J.R. Kirsch, Effects of anesthesia on cerebral blood flow, metabolism, and neuroprotection. J. Cereb. Blood. Flow. Metab. 38, 2192–2208 (2018).

45. P. Shin, W. Choi, J. Joo, W.-Y. Oh, Quantitative hemodynamic analysis of cerebral blood flow and neurovascular coupling using optical coherence tomography angiography. J. Cereb. Blood. Flow. Metab. 39, 1983–1994 (2019).

46. E. Moult, et al., Ultrahigh-speed swept-source OCT angiography in exudative AMD. Ophthalmic. Surg. Lasers. Imaging. Retina. 45, 496–505 (2014).

47. R. Leitgeb, L. Schmetterer, W. Drexler, A. Fercher, R. Zawadzki, T. Bajraszewski, Realtime assessment of retinal blood flow with ultrafast acquisition by color Doppler Fourier domain optical coherence tomography. Opt. Express. 11, 3116–3121 (2003).

48. I. Arganda-Carreras, et al., Trainable Weka Segmentation: a machine learning tool for microscopy pixel classification. Bioinformatics. 33, 2424–2426 (2017).

49. J. Schindelin, et al., Fiji: an open-source platform for biological-image analysis. Nat. Methods. 9, 676–682 (2012).

50. N. de Riva, et al., Transcranial Doppler pulsatility index: what it is and what it isn’t. Neurocrit. Care. 17, 58–66 (2012).

