## Supplementary Materials for "Increased capillary stalling is associated with endothelial glycocalyx loss in subcortical vascular dementia"

* Wang-Yuhl Oh

* Yong Jeong


**This PDF file includes:**

Figures S1 to S4

Legends for Movies S1

**Other supplementary materials for this manuscript include the following:**

Movies S1

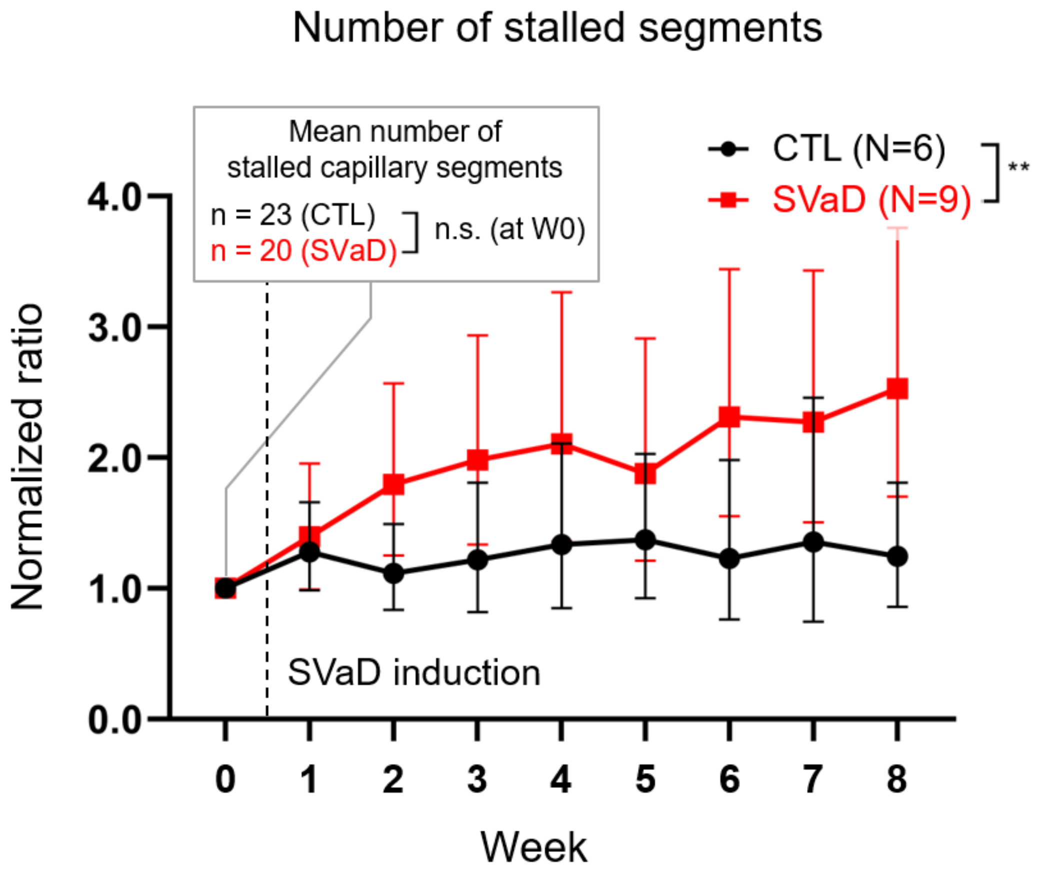


**Fig. S1. Changes in the number of stalled segments after inducing SVaD.** Changes in the number of stalled capillaries in control and SVaD mice were significantly different (control, n = 6; SVaD, n = 9; $F_{(8,104)} = 3.148$; p = 0.0031). Values are normalized to baseline. Log transformation was performed before statistical testing. Error bars represent the 95% confidence interval of the mean. The mean number of stalled capillary segments at W0 is depicted in the inlet box; no significant difference was found between control and SVaD mice (control, n = 6; SVaD n = 9; p = 0.4939). Two-tailed unpaired student t-test was used. Two-way repeated measure ANOVA test followed by Bonferroni's post-hoc was used (**: p<0.01 for interaction effect of group and time). SVaD, subcortical vascular dementia; ANOVA, analysis of variance.

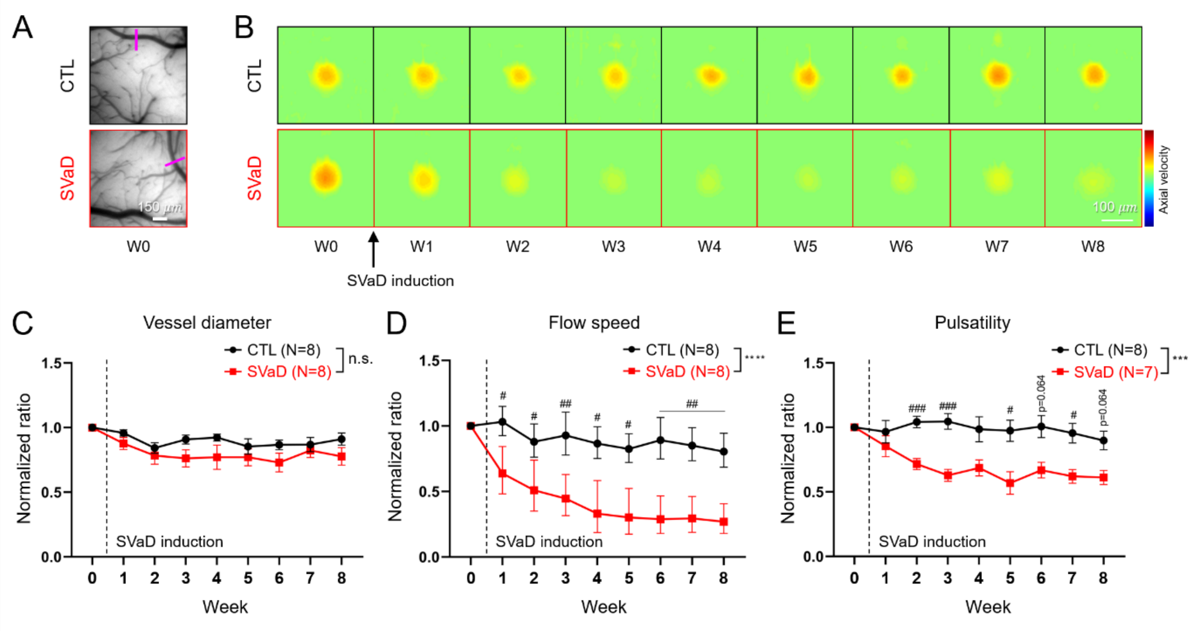


**Fig. S2. Changes in arterial blood supply after inducing SVaD.** A. CCD images showing the cranial window of control and SVaD mice. A magenta line in each image indicates the location where D-OCT imaging was performed. B. Doppler OCT images of pial arteries in control and SVaD mice showing an apparent decrease in arterial flow speed in SVaD mice. C. Changes in the diameters of pial arteries in control and SVaD mice were not significantly different (control, n = 8; SVaD, n = 8; $F_{(8,112)}$ = 1.179; p = 0.3178). Data are presented as mean ± s.e.m. D. Changes in the flow speed of pial arteries in control and SVaD mice were significantly different (control, n = 8; SVaD, n = 8; $F_{(8,112)}$ = 8.063; p < 0.0001, after log transformation). Error bars represent the 95% confidence interval of the mean. E. Changes in arterial pulsatility in control and SVaD mice were significantly different (control, n= 8; SVaD, n = 7; $F_{(8, 104)}$ = 3.903; p = 0.0004). Data are presented as mean ± s.e.m. All values are normalized to the baseline values. Two-way repeated measure ANOVA test followed by Bonferroni's post-hoc test was used (***: p<0.001, ****: p<0.0001 for the interaction effect of group and time, #: p<0.05, ##: p<0.01, ###: p<0.001 for multiple t test with Bonferroni's post-hoc test). SVaD, subcortical vascular dementia; CCD, charge-coupled device; OCT, optical coherence tomography; D-OCT, Doppler optical coherence tomography; ANOVA, analysis of variance.

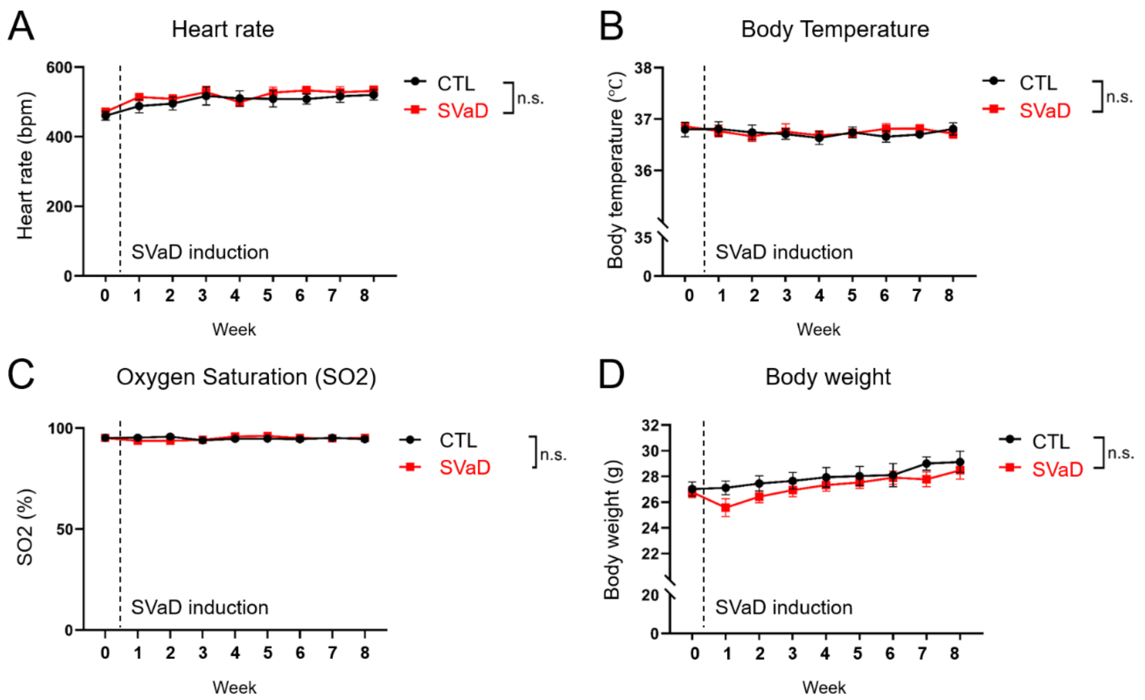


**Fig. S3. Physiological parameters in control and SVaD.** Changes in the heart rate, body temperature, arterial oxygen saturation, and body weight of control (n = 8) and SVaD (n = 9) mice are shown from A to D (heart rate: $F_{(8,104)} = 0.401$, $p = 0.918$, body temperature: $F_{(8, 104)}$, $p = 0.354$, arterial oxygen saturation: $F_{(8,104)} = 1.805$, $p = 0.084$, body weight: $F_{(8,104)} = 1.431$, $p = 0.192$). Data are presented as mean ± s.e.m. Two-way repeated measure ANOVA test followed by Bonferroni's post hoc was used. SVaD, subcortical vascular dementia; ANOVA, analysis of variance.

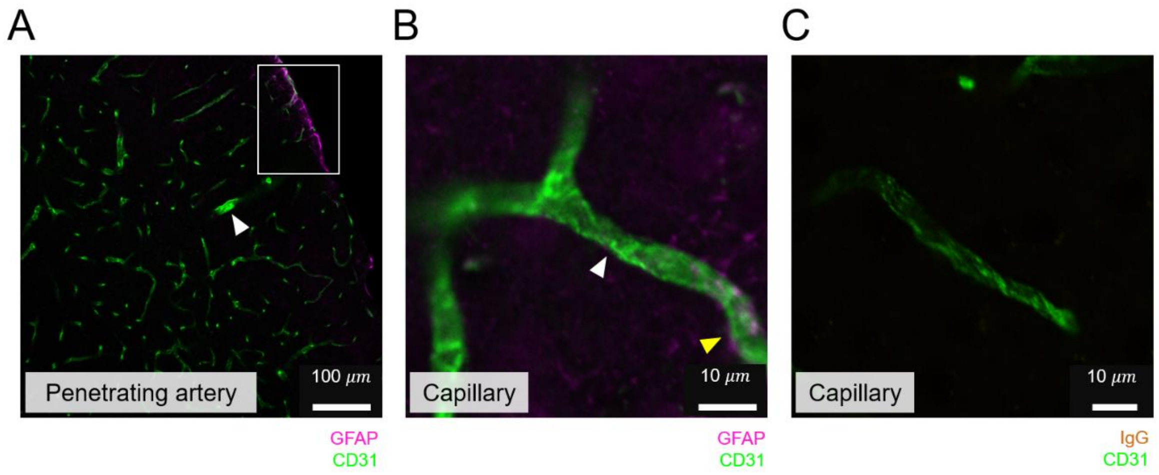


**Fig. S4. Immunohistochemistry images of control mice.** A. Reactive astrocytes were rarely observed around most vessels, including the penetrating artery (white arrowhead). Only a few reactive astrocytes and their endfeet were observed at the cortical surface and some of penetrating arteries, as shown in the white square. B. A capillary segment with no reactive astrocyte endfeet (white arrowhead) and with a small portion of astrocyte endfeet (yellow arrowhead). C. Capillaries without endogenous IgG leakage.

**Movie S1 (separate file). A video showing a temporal series of OCT angiograms acquired for 91 seconds.** Each angiogram was acquired in 7 seconds. Several regions indicated by yellow squares are selected and enlarged as shown on the right side of the video. Stalled capillaries are indicated by yellow arrows in each region. OCT, optical coherence tomography.
